# Drug-Eluting, Radiopaque, Tumor-Casting Hydrogels for Endovascular Locoregional Therapy of Hepatocellular Carcinoma

**DOI:** 10.1101/2025.11.25.690505

**Authors:** Yuxi C. Dong, Kathleen E. Villaseñor, Seokyoung Yoon, Ariful Islam, Luis Vazquez, Alexey Gurevich, Shaun McLaughlin, Terence P. Gade, David P. Cormode

## Abstract

Hepatocellular carcinoma (HCC) is one of the most common cancers worldwide. For patients with unresectable disease, locoregional therapies, such as transarterial chemoembolization (TACE), are the standard of care. However, even with this treatment, the average survival rate is only 27% at 5 years. Recent studies have demonstrated the potential of novel approaches to improve responses through more complete embolization and targeting ischemia-induced molecular dependencies. To build on these findings, we developed an injectable chitosan-based hydrogel system (AuNP-Lys05-gel) that we hypothesized might be a more effective embolic material by (i) enabling a more complete filling of tumor feeding blood vessels in a cast-like manner, (ii) encapsulating Lys05, a potent autophagy inhibitor, and (iii) providing X-ray contrast for direct visualization of the embolization. Chitosan hydrogels of varying composition were formed and characterized. We found that Lys05 could be loaded in this type of hydrogel and that it was released gradually over 7 days. We found that AuNP was better retained by the hydrogel than iopamidol, an FDA-approved contrast agent, indicating that AuNP would provide longer-lasting CT monitoring. In a rat model of HCC, AuNP-Lys05-gel reduced tumor growth and resulted in a 67% objective response rate vs 26% for a clinically used particle embolic. Histology indicated effective vascular occlusion by the gel. These findings indicate that AuNP-Lys05-gel merits further investigation as a treatment for HCC.

## Introduction

HCC is the most common primary malignancy of the liver, is also one of the most incident cancers, and the most common cause of cancer-related death worldwide.[1, 2] It is estimated that there are over 500,000 new cases per year, globally.[3, 4] Among numerous underlying causes that induce HCC, cirrhosis remains the most important risk factor for the development of HCC with more than 80% of individuals diagnosed with HCC having preexisting cirrhosis.[4] Hepatitis B and C, as well as alcohol consumption, diabetes and high-fat diets are causes of cirrhosis.[5] Despite the overall decreasing mortality rates for cancer, mortality for HCC continues to increase in men and women, owing to lifestyle factors that lead to cirrhosis, the challenges of early diagnosis of HCC, and the consequent late stage of diagnosis.[6, 7] There are numerous options for the treatment of patients with HCC, including liver transplantation, radiofrequency ablation, trans-arterial embolization (TAE), transarterial chemoembolization (TACE), radioembolization, as well as systemic targeted agents such as immunotherapy.[8] The decision for the treatment is multifactorial, including the disease burden in combination with patient functional status, and is usually discussed at a tumor board amongst many subspecialists.

Among various treatment options, TACE is the most commonly used therapy for HCC worldwide.[4, 9] TACE is an image-guided treatment performed by interventional radiologists targeting tumor-feeding vessels to restrict the blood supply to a tumor in the liver while also delivering chemotherapeutics. TACE is performed by delivering the chemotherapeutics and embolics directly into the tumor in the liver using a microcatheter.[10] Conventional TACE (cTACE) system includes the administration of a drug-in-oil emulsion using Lipiodol followed by an embolic material.[11] Despite proven efficacy, recurrence following TACE is common and results from limitations in the ability to achieve durable embolization as well as to optimize delivery of targeted therapeutics. While preclinical and clinical studies demonstrate that TACE enables the concentrated delivery of therapeutic agents, achieving tissue concentrations that are an order of magnitude greater than those achievable through systemic delivery with tumor-to-normal parenchyma ratios ranging from 3:1 to 20:1, there is usually a peak of chemotherapeutic agent in the bloodstream observed immediately after the TACE procedure, which can lead to systemic side effects.[3, 12–17] The drug-eluting bead (DEB) for TACE was therefore developed to integrate the delivery of the chemotherapeutic with the embolic. However, the large size of these beads compared to the size of tumor capillaries (the beads are 100-300 µm[2] whereas the capillaries measure up to 30 µm in diameter) may cause incomplete embolization. Released drug may therefore not diffuse deep into the tumor, leading to reduced therapeutic efficacy.[2, 18–20] At the same time, most beads are radiolucent and cannot be seen with any form of imaging modality, while the radiopaque beads are challenging to visualize,[21] making it difficult for interventional radiologists to locate them and determine adequate delivery to, as well as distribution within target tumors. Therefore, an optimal chemoembolization platform for HCC treatment requires the following three features: 1) an efficient delivery vehicle tailored for locoregional delivery; 2) mechanical flexibility to allow occlusion down to the capillary level; and 3) direct visualization under fluoroscopic guidance (fluoroscopy is an X-ray based imaging technique that is a mainstay of interventional radiology therapeutic procedures, including TACE).

In this study, we developed chitosan-based thermosensitive, radiopaque hydrogels that can be loaded with targeted agents. Lys05, an autophagy inhibitor, is selected as the loading drug since it was recently demonstrated that it effectively targets HCC cells that survive TACE-induced ischemia.[22] A hydrogel material should exhibit several characteristics in order to be effectively used in this clinical application, including the ability to maintain its integrity for a desired period of time, as well as good biocompatibility and biodegradability.[23, 24] Chitosan-based hydrogels are known to be biocompatible and capable of degradation via human enzymes.[25] The system that we propose to use is liquid at room temperature, but when injected into the body and reaches body temperature, the material undergoes a rapid transition to a hydrogel,[26, 27] which provides its potential for use as a liquid embolic. Upon reaching the tumor arterioles and capillary beds, where blood flow is relatively low, the liquid gel will quickly fill the blood vessels, then rapidly form a solid cast within the blood vessels, entirely blocking blood flow and providing a drug depot throughout the tumor, to facilitate maximal tumor killing via ischemia and drug delivery. Importantly, the material will be used to encapsulate AuNP, which will allow the interventional radiologist to directly visualize the material during the TACE procedure to assess the completeness of target tissue embolization, thereby enhancing treatment efficacy. In addition, the AuNP will allow the hydrogel to be monitored over time via follow-up CT. We herein report the characteristics of this hydrogel, *in vitro* HCC cell killing, and *in vivo* tumor treatment effects.

## Results

### Formation and characterization of hydrogels

The thermosensitive hydrogels were formed by ionically crosslinking chitosan and ammonium sodium hydrogen phosphate (AHP) using a modification of a previously reported method.[28] To encapsulate AuNP and Lys05 in the hydrogels, the desired cargoes were mixed with AHP first before reacting with chitosan for crosslinking. Sub-5 nm AuNP were used as the renal filtration cut-off threshold is about 6 nm.[29, 30] Renal clearance of nanoparticles is desirable to facilitate clinical translation of the hydrogel. The sol-gel transition was initiated when the temperature was changed from room temperature to 37 °C, i.e. body temperature. The formation of the hydrogel was indicated by passing the vial inversion test (i.e., the gel stops flowing when the vial containing it is inverted) (**Figure 1A**). A transmission electron micrograph of the resulting hydrogels is shown in **Figure 1B**, where the AuNP dispersion in the hydrogel was confirmed.

**Figure 1.**
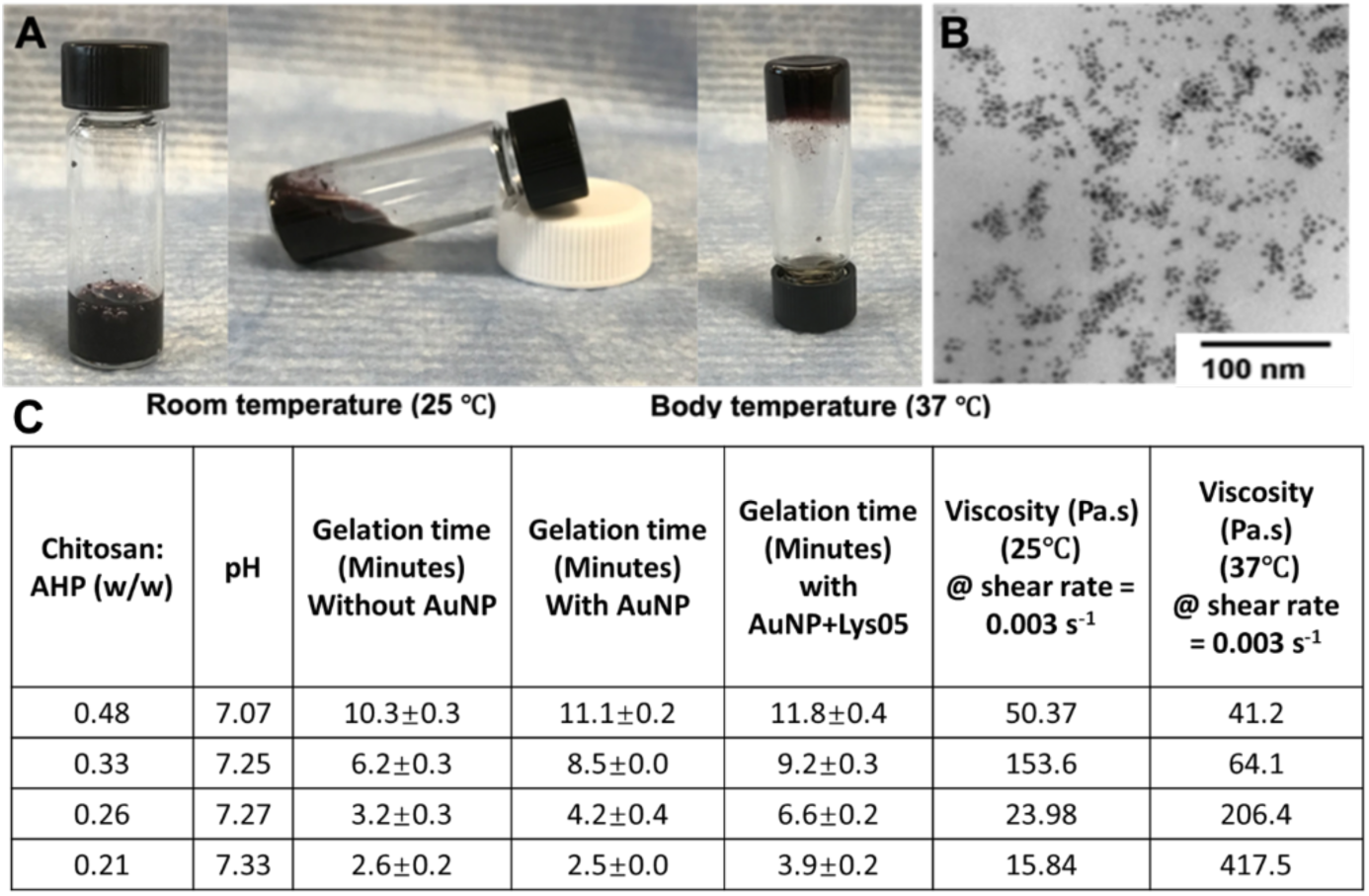
Characterizations of thermosensitive hydrogels. **(A)** Photos of thermosensitive hydrogels loaded with AuNP and Lys05. **(B)** TEM micrograph of sub-5 nm AuNP dispersed in the hydrogel. **(C)** Hydrogels with different polymer/crosslinker mass ratio and their corresponding pH, gelation time and viscosity.

Interestingly, by varying the chitosan to AHP mass ratio, the gelation time of the hydrogels can be tuned. Higher chitosan to AHP mass ratios resulted in longer gelation times (**Figure 1C**). The encapsulation of AuNP prolonged the gelation time, possibly due to the dilution of crosslinker AHP with the addition of AuNP. In addition, the pH of the corresponding hydrogel solutions varied from 7.07 to 7.33. The viscosity and storage modulus of the hydrogel solution was found to increase after the increase of temperature from 25 °C to 37 °C, further confirming the gelation of the hydrogels when they reached body temperature (**Figure 2**). The mass ratio of 0.21 was used for all the subsequent experiments, as the resulting hydrogels have a sufficiently low viscosity so that they can be injected through a microcatheter (i.e., MTV-40 tail vein catheter), but will also gel within four minutes, such that they will quickly gel upon deployment in the patient.

**Figure 2.**
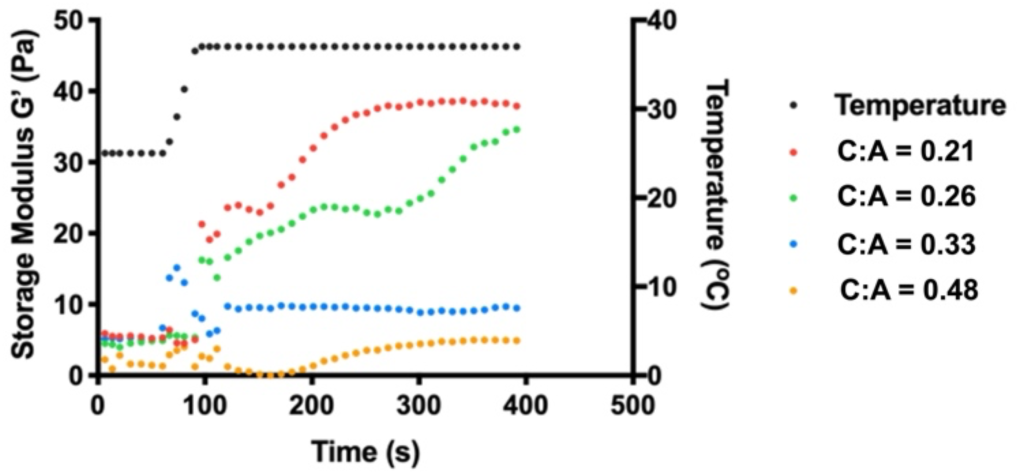
Rheological characterization of hydrogels. Temperature-dependent storage modulus of hydrogels with different polymer to crosslinker mass ratios measured using oscillatory shear rheometry.

### Hydrogel degradation and drug release

The cumulative release of AuNP and Lys05 from the hydrogel was investigated *in vitro*, through incubations in FBS-supplemented PBS. Small amounts of supernatants were taken out at different time intervals and analyzed for both AuNP and Lys05 concentrations by spectrophotometry using their respective characteristic absorbance peaks (i.e., 328 and 520 nm). As can be seen in **Figure 3A**, there was a loading dependency of AuNP release (i.e. greater release was observed with higher AuNP loading). However, there was only 11% of the payload released even at the highest loading. On the other hand, 34% of an iodinated contrast agent was released over the same period. The release of AuNP was significantly lower than that of the iodinated agent, possibly due to the larger size of the AuNP compared to the iodine molecules. High retention of contrast agents in the hydrogel is desired so that the hydrogel can be monitored post-embolization through CT imaging; therefore, AuNP are advantageous for this purpose. As indicated in Figure **3B**, Lys05 has somewhat higher release kinetics compared to DOX, possibly since DOX is a larger molecule. Moreover, there was no significant difference in release between the drug loadings tested, indicating that the loading doesn’t affect the drug release kinetics, within the tested range.

**Figure 3.**
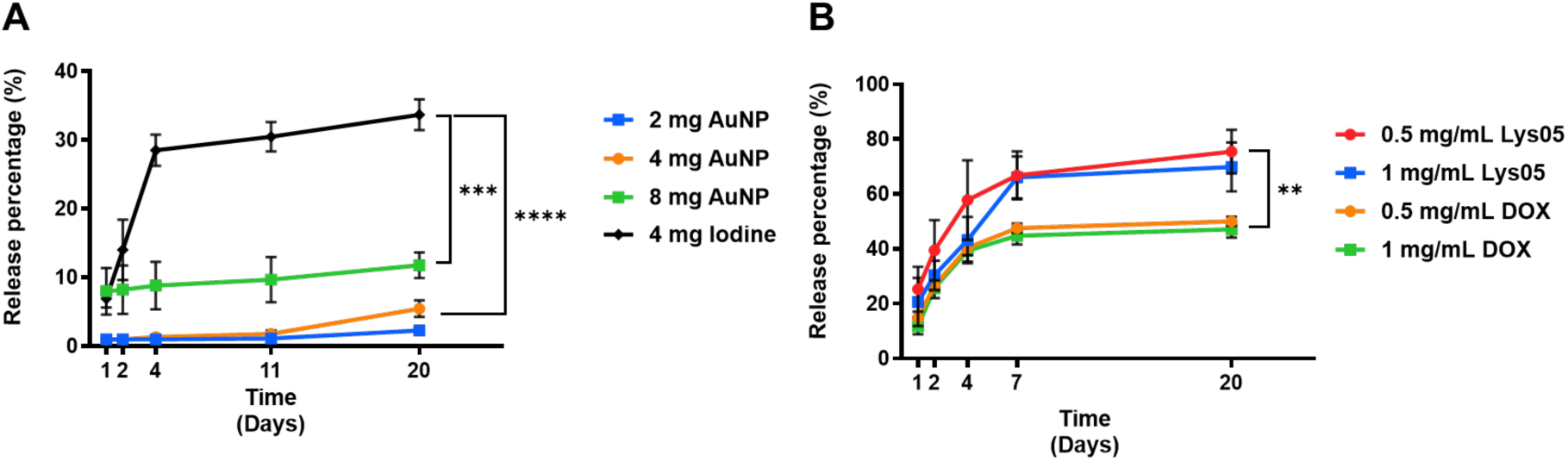
Release profiles of AuNP, iodine molecules, Lys05, and DOX from hydrogels. Release profiles of **(A)** AuNP and iodinated molecules, as well as **(B)** Lys05 and DOX from hydrogels over 20 days.

### Contrast generation of hydrogels

Next, we performed phantom imaging to examine the contrast generation of AuNP-Lys05 hydrogels with a small animal microCT imaging system (**Figure 4A**). A linear correlation of AuNP concentration in the hydrogels with attenuation was observed (**Figure 4B**). Moreover, even at loadings as low as 2 mg Au/ml, the contrast generated was >100 HU. This result supports our hypothesis that the hydrogel can be monitored through CT imaging.

**Figure 4.**
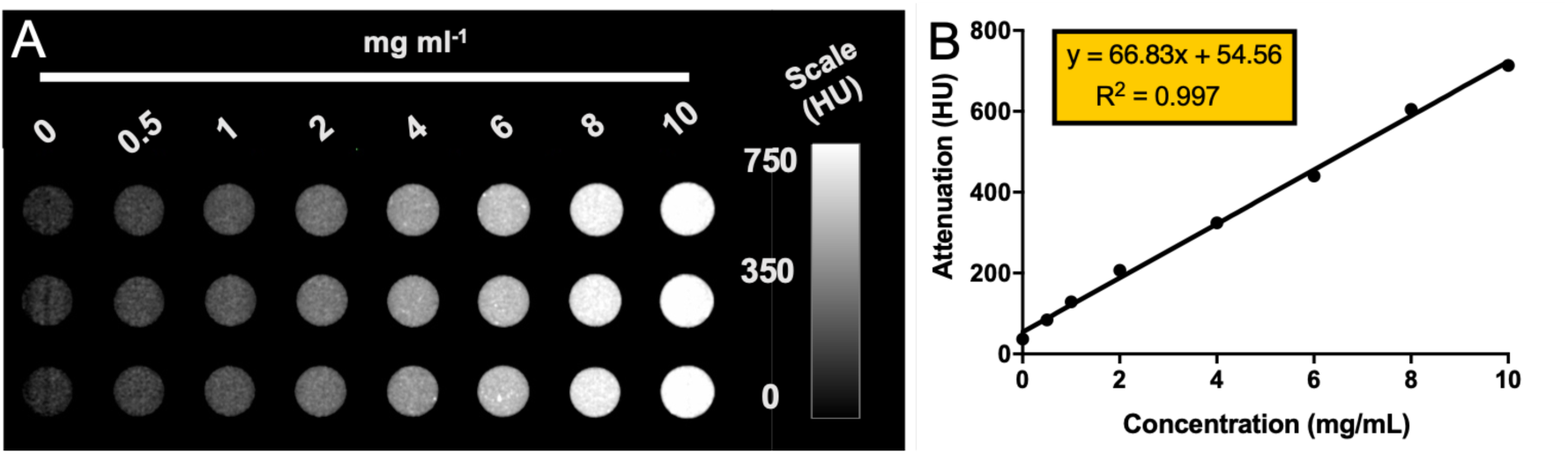
Phantom imaging of AuNP-Lys05 hydrogels with CT. **(A)** CT phantom image and **(B)** X-ray attenuation changes versus concentration for hydrogels scanned by a microCT system.

### *In vitro* cytocompatibility

To assess the biocompatibility of chitosan-based hydrogels containing AuNP contrast agents, hydrogels loaded with 8 mg/mL AuNP were incubated with DMEM cell medium for various durations (i.e., 1, 2, 7 days). The supernatants containing hydrogel degradation byproducts and AuNP were incubated with HCC cells (Huh7), macrophages (J774A.1) and endothelial cells (SVEC), as those are the cell types that would potentially interact the most with the hydrogel *in vivo*. Since we propose to treat HCC with hydrogels, we selected the Huh7 cell line as a clinically relevant model of HCC cancer. The effects on cell viability were measured using the LIVE/DEAD assay. As shown in **Figure 5A**, incubation with hydrogel degradation byproducts and released AuNP did not affect cell viability, underscoring the safety of the AuNP used in the drug-loaded hydrogel and the non-drug loaded hydrogel. The cancer-killing effects of AuNP-Lys05 hydrogel were then assessed on Huh7 HCC cells in relevant microenvironments – standard conditions (21% oxygen) or hypoxic conditions (1% oxygen). Lys05- or DOX-loaded hydrogels were incubated with cell media for 48 hours to allow for drug release. Then, the supernatants were collected at different time points and used to treat the cells, after which the MTS assay was performed. For cells treated with released Lys05, the hypoxic condition resulted in significantly lower cell viability compared to the standard condition. However, this difference was not observed in cells treated with DOX, moreover, the reductions in viability of Huh7 cells treated with DOX was modest. The results confirmed the previously reported finding that Lys05 is more effective under ischemia (**Figure 5B**).[31]

**Figure 5.**
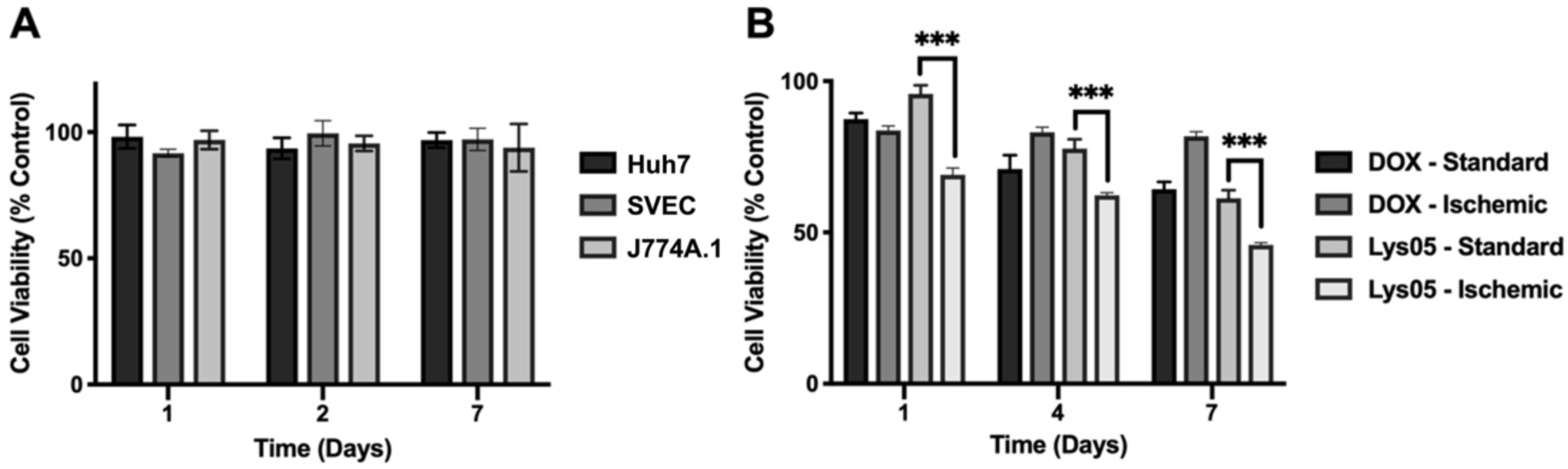
*In vitro* cytocompatibility and cytotoxicity of AuNP-Lys05 hydrogels. **(A)** Viability of cells incubated with hydrogel degradation byproducts and released AuNP. **(B)** Viability of Huh 7 cells after receiving treatment with drugs eluted from hydrogels.

### In vivo treatment of HCC in a rat model

Next, we performed an *in vivo* study using a diethylnitrosamine (DEN)-induced autochthonous rat model of HCC.[32, 33] These animals were embolized with either AuNP-Lys05-gel (8 mg/ml AuNP and 1 mg/ml Lys05 loading) or a particle embolic (PE, Embozene, Varian) or were administered a sham treatment. Following treatment, tumor growth was monitored by serial MRI. As can be observed from representative images (**Fig. 6A**) and plots of tumor growth over time (**Fig. S1,2**), AuNP-Lys05-gel slowed tumor growth compared to sham or PE treatment. In addition, the administered AuNP-Lys05-gel could be clearly visualized on CT scans (**Fig. 6B&C**). Response Evaluation Criteria in Solid Tumors (RECIST)[34] were used to assess the response of the tumors to treatment (**Fig. 6D-G**). We found that AuNP-Lys05-gel resulted in substantially improved objective response rates (67%) as compared to either sham (where no objective responses were observed) or PEs (26%, p=0.038 via Pearson’s Chi-squared test with Yates’ continuity correction).

**Figure 6.**
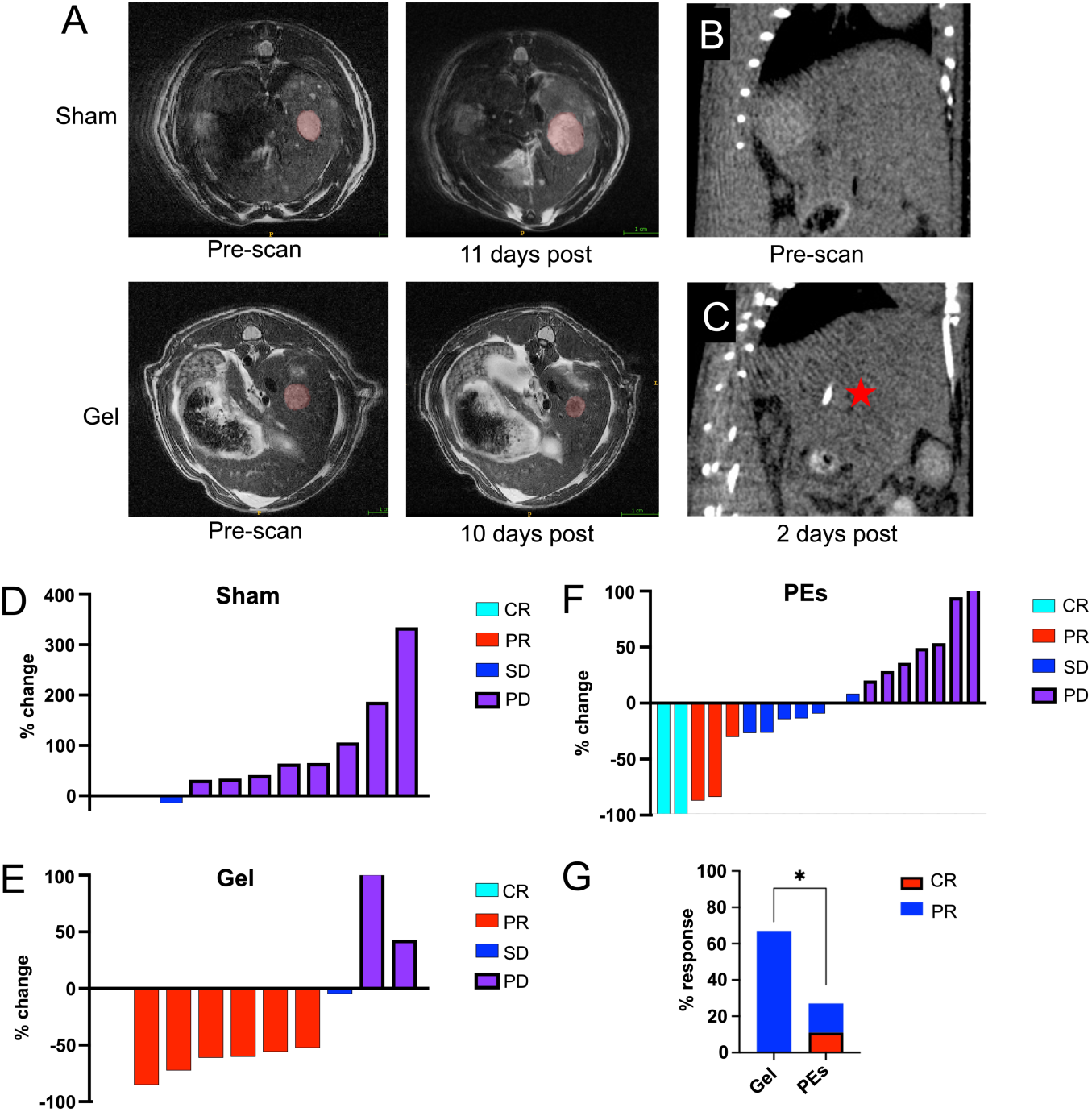
A) Representative MR images of liver tumors treated with either sham or AuNP-Lys05-gel (‘Gel’) in the DEN rat model. CT scans of the liver of a rat B) before and C) two days post administration of AuNP-Lys05-gel (indicated by red star) into an HCC tumor. RECIST criteria responses of tumors to D) sham, E) AuNP-Lys05-gel and F) PE treatment. CR = complete response, PR = partial response, SD = stable disease and PD = progressive disease. G) Objective response rate of tumor treated with AuNP-Lys05-gel or PEs (* = p<0.05).

### Penetration of tumors with hydrogel

Excised tumors that were either embolized with AuNP-Lys05-gel or underwent a sham procedure were prepared for histology and stained for chitosan as described previously (**Fig. 7A**).[35] Substantial penetration of the tumor vasculature was observed with visualized penetration of vessels down to 50 μm, supporting the concept that this hydrogel would result in extensive embolization and resultant ischemia. It is also noteworthy that treated tumors were highly necrotic (e.g. lack of the blue nuclear stain) compared to sham treated. In addition, electron microscopy was performed on tumor tissue (**Fig. 7B**), where gold nanoparticle deposits were observed within the tissue.

**Figure 7.**
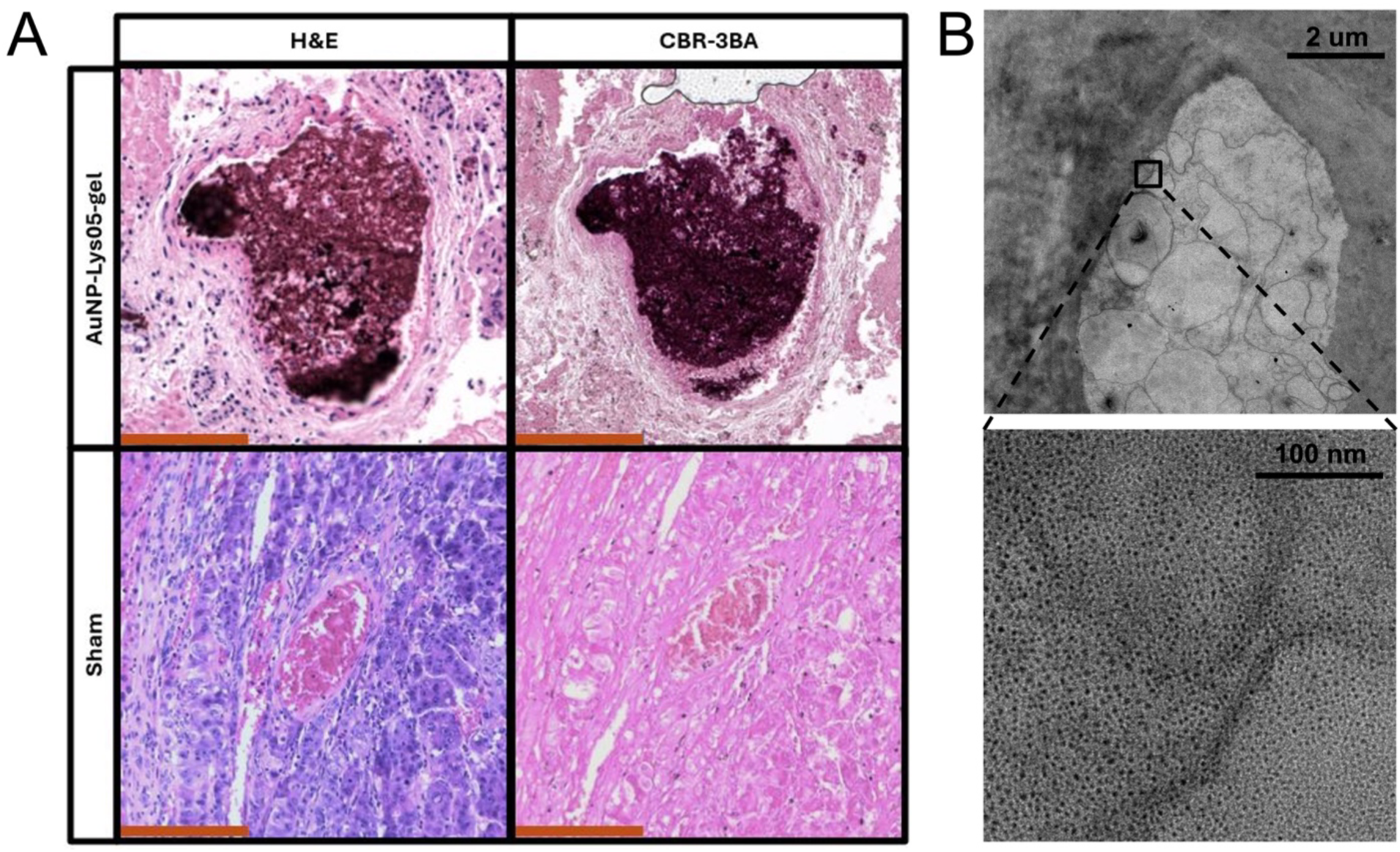
**(A)** Histology of tumors that were embolized either with AuNP-Lys05-gel or underwent a sham procedure. Stains are hematoxylin and eosin (H&E) and Cibacron Brilliant Red-3BA (CBR-3BA). **(B)** Electron microscopy of tumor tissue embolized with AuNP-Lys05-gel.

## Discussion

In this study, we have developed a radiopaque, thermosensitive hydrogel as an embolic for endovascular locoregional therapy. The hydrogel is liquid at room temperature but undergoes a rapid transition to a hydrogel once it reaches body temperature. The presented data demonstrate the ability of the liquid hydrogel to fill the blood vessels within the HCCs, blocking the tumor blood flow and providing a drug depot throughout the tumor. Chitosan-based hydrogels can persist for periods of months, ensuring a durable period of hypoxia.[36] Moreover, chitosan-based hydrogels can gel at low polymer concentrations, while alternatives such as Pluronics are non-biodegradable, gel at high concentrations (e.g., 20%), and dissolve in aqueous media rapidly.[37] In fact, the development of chitosan hydrogels has drawn intense interest based on its potential use in various biomedical applications.[38–41] Due to its thermogelling properties when crosslinking with polyol counterionic dibase salts such as α-glycerophosphate (GP) or AHP, thermosensitive chitosan hydrogels are also highly attractive as drug carriers.[42–44] Herein, we used thermosensitive chitosan hydrogels as a “two-in-one” platform for both therapeutic delivery and embolization.

Several hydrogel formulations have been explored for this application. For instance, temperature-sensitive p(N-isopropylacrylamide-co-butyl methylacrylate) nanogels, chitosan/glycerophosphate hydrogels, sodium alginate-derived hydrogels have been studied for interventional embolization treatment.[45–47] Other “smart” hydrogels have been studied for the same purpose as well. For instance, Nguyen et al. used a pH-sensitive anionic block copolymer comprised of poly(ɛ-caprolactone-co-lactide) (PCLA), poly(ethylene glycol) (PEG) and poly(urethane sulfide sulfamethazine) (PUSSM) as a DOX-containing embolic agent for chemoembolization for HCC. The polymer existed as a liquid at high pH but transitioned to hydrogels when injected into the extracellular tissues of the tumor site, where the microenvironment is more acidic.[11] Similarly, work performed by Lym et al. demonstrated intra-arterial administration of pH-sensitive hydrogels in a rabbit VX2 liver tumor model.[48] However, few of these hydrogels are radiopaque, presenting challenges to interventional radiologists when the assessment of blood flow and deployment of hydrogels are still needed. Although TACE procedures typically include the mixing of iodinated contrast agents with embolics to visualize changes in vascular flow, the eventual location of microspheres cannot be detected, making it difficult for interventional radiologists to judge whether there is sufficient delivery of embolic agents to the tumor site.

Our work demonstrated the contrast generation of hydrogels *in vitro* and *in vivo*. The radiopacity of hydrogels was enabled by the loading of contrast agents AuNP. The encapsulation of AuNP not only allows the deployment and retention of the hydrogel to be assessed over time through serial imaging with CT, but also enables the K-edge imaging feature of spectral photon counting CT (SPCCT) for future ‘hot-spot’ imaging of the hydrogel biodistribution, allowing the gel to be distinguished from iodinated agents used to image blood flow.[49, 50]

The current study has several limitations. The safety and *in vivo* biodegradation/excretion of the hydrogels remains to be studied, although the safety is expected to be favorable, since chitosan has the FDA designation of ‘Generally Regarded As Safe’ (GRAS),[51, 52] and similar AuNP-loaded hydrogels had good safety profiles.[53, 54] In addition, while the *in vivo* experiments demonstrated the efficacy of the AuNP-Lys05-gel, further experiments will be needed to determine the relative contributions of the thermosetting embolic materials versus the release of the drug. Finally, while the DEN rat model of HCC has been demonstrated to recapitulate human disease, further studies are required to determine the effectiveness of the AuNP-Lys05-gel for targeting human HCC *in vivo*.

## Conclusion

In summary, we report the development of an injectable, radiopaque, drug loaded thermosensitive hydrogels that rapidly thermosets at body temperature for use in endovascular locoregional therapy. We have demonstrated excellent gelation time control of the hydrogel, efficient loading of contrast agents and drug payloads, and favorable drug release profiles *in vitro*. We found that the autophagy targeting drug, Lys05 showed a marked increase in cell anti-proliferative activities after exposure to ischemic conditions. In addition, *in vitro* CT imaging results confirmed the robust x-ray contrast generated by the encapsulated AuNP. Our *in vivo* findings point to the AuNP-Lys05-gel being a more effective treatment for HCC as compared to embolization alone, meriting further investigation.

## Materials and methods

### Materials

Gold (III) chloride trihydrate (99.9%), sodium borohydride powder (98.0%), L-glutathione reduced (GSH), chitosan (medium molecular weight), ammonium sodium hydrogen phosphate (AHP, NaNH_4_HPO_4_·4H_2_O), and doxorubicin hydrochloride (DOX) were purchased from Sigma-Aldrich (St Louis, MO). Lys05 was purchased from Selleck Chemicals (Houston, TX). HepG2, Renca, and SVEC4-10 cell lines were purchased from ATCC (Manassas, VA). LIVE/DEAD assay kits were purchased from Life Technologies Invitrogen (Grand Island, NY). Cells were cultured in Dulbecco’s Modified Eagle Medium supplemented with 10% fetal bovine serum and 1% penicillin (10000 units/mL) from Life Technologies Invitrogen (Grand Island, NY). Milli-Q deionized water (18.2 MΩ cm) was used throughout the experiments.

### AuNP synthesis and characterization

AuNP were synthesized via the following method. In brief, gold (III) chloride was reduced with sodium borohydride and subsequently modified with glutathione as a stabilizing coating ligand.[55] The nanoparticles were purified with DI water and concentrated to a final volume of 1 mL for use in subsequent experiments. ICP-OES was performed to determine the final concentration of the stock AuNP solution. The core size of AuNP was examined with TEM using a JEOL 1010 microscope (JEOL USA Inc., Peabody, MA) at 80 kV. The UV-vis absorption spectra of diluted AuNP were recorded with a Genesys UV-visible spectrophotometer (Thermo Scientific, Waltham, MA).

### Thermosensitive hydrogel formation

The procedure to form chitosan-based thermosensitive hydrogel was adapted from previous studies.[28, 56] To prepare the 2% w/v chitosan solution, 2.0 g chitosan was dissolved in 100 mL of 0.75% acetic acid under magnetic stirring for 24 hours at room temperature. 60% AHP solution was prepared by dissolving 6 g of AHP powder in 10 mL of DI water. Varying amounts of AHP solution (i.e., 70, 90, 130 or 160 μL) were quickly added to 1 mL of chitosan solution in an ice bath and vortexed for 30 seconds until the solution turned turbid. To prepare Lys05-AuNP loaded hydrogel solutions, 100 μL of 0.5-5 mg/mL Lys05 was added to 125 μL of 64 mg/mL AuNP solution and then mixed with AHP solution, and subsequently mixed with chilled chitosan solution as described above. The liquid hydrogel solution was stored at 5 °C before gelling. To complete the sol-gel transition, the chitosan-AHP mixture was incubated in a water bath at 37 °C. The gel formation was confirmed by assessing the flowability of the solution when the vial was inverted.

### Gelation time determination

To determine the effect of chitosan-to-AHP ratio on the time of sol-gel transition, different amounts of AHP (100, 120, 140 or 160 μL) were added to 1 mL of 2.5% w/v chitosan solution in an ice bath. The vial inversion method was used to determine the time needed for the chitosan-AHP mixture to gel. The mixtures were incubated in a water bath at 37 °C with the test vials fully immersed in water. The vials were taken out every 30 seconds from the water to quickly check the flowability of the solution by inverting the vials. The time at which the solution stopped flowing was taken as the gelling time. The gelling times were reported as the average of four tests for each formulation.

### Rheological analysis and injectability

The rheological properties and injection force needed for the non-gelled solutions and the hydrogels were measured using a TA Instruments RFS3 rheometer and a Tekscan Force Sensor, respectively, as previously published.[57] The hydrogels were loaded onto the Peltier plate on a rheometer. A parallel plate geometry with a gap size of 100 μm was used. The following measuring conditions were used for the measurements: frequency sweep (0.001-500% strain, 10 Hz), continuous flow (shear rates from 0 to 50 s^-1^ over 2 minutes 30 seconds), and cyclic strain (low = 0.2% strain, high = 500% strain, 10 Hz). The measurements were carried out at 25 °C from 0 to 150 seconds and at 37 °C from 150 to 300 seconds.

To measure the injection force of the non-gelled solutions, 500 μL of the sample were loaded into a 1 mL syringe assembled with an Excelsior SL-10 microcatheter (Stryker, MI) or a rat tail vein catheter (SAI Infusion Technologies, IL). The syringe was placed into an Instron 5848 MicroTester (Norwood, MA. The experiment was conducted with a flow rate of 2 mL/hr and the maximum force applied during hydrogel extrusion over time was recorded. Testing was carried out in air and repeated 4 times.

#### Transmission Electron Microscopy

AuNP loaded hydrogel samples were fixed with 2% paraformaldehyde and 2.5% glutaraldehyde, embedded, cut into sections of 60 nm in thickness and mounted onto copper grids. A JEOL 1010 microscope (JEOL, Tokyo, Japan) was used to acquire TEM images of hydrogels at 80 kV.

#### Drug and AuNP release studies

*In vitro* release studies were performed to determine the percentage cumulative release of Lys05 and AuNP from the hydrogel over time. Hydrogels loaded with DOX and iodinated contrast agent were used as control groups for comparison. 0.5 or 1 mg of drug and 1, 2, 4 or 8 mg of AuNP were added to 1 mL of chitosan-AHP mixture and gelled at physiological temperature (37 °C). The resulting hydrogels were placed in a 100-fold higher volume of 10% FBS in PBS at 37 °C in a water bath. 1 mL of supernatants were taken from the larger volume after 1, 2, 4, 7, and 20 days of incubation and analyzed for both drug and contrast agent concentrations. The gold and iodine content were determined by ICP-MS, and the Lys05 and DOX content were analyzed by spectrophotometry using their respective characteristic absorption peaks (i.e. 328 nm and 480 nm, respectively).

#### CT phantom imaging

The X-ray contrast generation of the hydrogels was investigated using a micro CT system. Samples were scanned in custom made plastic phantoms to simulate the effect of beam hardening experienced within subjects. Samples were prepared in triplicate, and a range of concentrations (0 – 10 mg/mL) were scanned using an MILabs micro CT scanner at a tube voltage of 55 kV and a tube current of 190 μA with an exposure time of 75 ms. Slices of 100 μm thickness with an increment of 0.1 mm were reconstructed using the algorithm provided by the manufacturer. Image analyses were performed using OsirixX (v.3.7.1 64-bit software). A circular ROI was drawn on the coronal view of each tube. The CT attenuation value for each sample tube at each concentration was recorded as the average of three samples of that same concentration.

### Cell culture

#### *In vitro* biocompatibility

The LIVE/DEAD assay was performed to assess the biocompatibility of non-drug loaded hydrogels. Hydrogels loaded with 8 mg/mL AuNP were incubated with DMEM cell medium for various durations (i.e., 1, 2, and 7 days). The hydrogel-treated cell medium was then incubated with Huh7 (HCC cell line), J774A.1 (macrophage cells), and SVEC4-10EHR1 (endothelial cells), as those are the cell types that would potentially interact the most with the hydrogels. After 24 hours of incubation at 37 °C and 5% CO_2_, the cells were incubated with LIVE/DEAD cocktail and imaged with a Nikon Eclipse Ti-U fluorescence microscope with FITC (ex: 495, em: 519 nm) and Texas Red (ex: 595, em: 613 nm) filters to image live and dead cells, respectively. Four different areas of each well were imaged. The number of live and dead cells was measured using a custom MATLAB (Mathworks, Natick, MA) code. The percentage viability was determined by calculating the ratio of live to dead cells normalized to control.

#### Effect of Lys05 release on cells

The MTS assay was performed to assess the cytotoxicity of the drug released from the hydrogel on HCC cells (i.e. Huh7) under standard and hypoxic conditions, by adaptation of previously reported procedures.[58] A previous study demonstrated that the autophagy targeting drug, Lys05 is more effective under ischemic conditions compared to doxorubicin.[31] For this experiment, media incubated with hydrogels loaded with either Lys05 and doxorubicin for 1, 4, and 7 days were used to treat Huh7 cells for 24 hours at both standard conditions (21% oxygen) and hypoxic conditions (1 % oxygen with ischemic media – 2% FBS, low glucose, low glutamine). Once the treatment was done, the cells were washed with DPBS and incubated with the MTS assay solution (Promega, Madison, WI) comprised of 20 uL of stock MTS solution and 100 uL of cell medium. After the 2-hour incubation period, a microplate reader (Synergy H1, BioTek, VT) was used to record the absorbance at 490 nm. The cell viability relative to control (%) for each concentration and treatment duration was determined.

#### Animal model

Male Wistar rats weighing 300–400 g (Charles River Laboratories) were used, adhering to a protocol approved by the Institutional Animal Care and Use Committee. The induction of HCC in the rat liver was accomplished through the ad libitum oral administration of 0.01% diethylnitrosamine over a 12-week period. Tumor diagnosis in rats was done via T_2_-weighted MRI scans (Agilent 4.7-T 40-cm horizontal bore MRI) at 5–7-day intervals starting from the 11th week of the diet. The tumor volume was measured using ITK-SNAP software. Once the tumor reached a volume within the range of 100-300 mm³, it was considered suitable for embolization.

#### Embolization and MRI tracking

Rats were anesthetized with isoflurane and then positioned on a draped surgical bed in a supine position, with continuous administration of 2% isoflurane through a nose cone. The limbs and tail were securely taped down, and the skin surface between the right hind leg and lower right abdomen was prepared for the surgical incision. A small incision was made to access the right common femoral artery (CFA). Tissues were manipulated using a self-retaining retraction system to provide access to the femoral artery. Under a stereomicroscope (Leica M80), the muscle and vascular sheath were meticulously dissected to isolate the femoral artery and an arteriotomy was created by a 26G needle in the right CFA.

Thereafter, the procedure involved fluoroscopy-guided intervention, including cannulation using a 1.7-F microcatheter, introduction of a guide wire, and the identification of hepatic arteries as well as tumor-bearing arteries. Prior to embolization, a digital subtraction angiography was performed using an Angiostar Plus or Cios imaging system (Siemens). This involved capturing detailed images to visualize the vascular structures and aid in the precise identification of hepatic arteries and tumor-bearing arteries. Tumors measuring at least 100 mm^3^ in volume were selected for the experiments and a volume ranging from 75 to 200 microliters (adjusted based on the tumor and vessel’s diameter) of AuNP-Lys05 hydrogel (made as noted above, loaded with 8 mg AuNP and 0.5 mg Lys05) was delivered to the tumor-feeding artery through selective catheterization. This process was carried out using the Standard PHD ULTRA™ CP Syringe Pump from Harvard Apparatus, at a controlled rate of 75 to 100 microliters per minute. Following this, an additional round of post-embolization arteriography was conducted to ensure the openness of vessels, excluding the target artery. Subsequently, the catheter was removed, the arteriotomy was closed, and the incised area was sutured using 4-0 sutures. Post-surgery, close observations were made on the rats’ recovery, encompassing aspects such as waking up, overall health, walking, and their consumption patterns.

All treated rats underwent weekly MRI screenings for up to 4 weeks, provided their health and condition remained good. Tumor volume was monitored to assess changes after treatment. Regular recordings were maintained for weight and other physical appearances. At the conclusion of the study rats was euthanized and tissues were collected for future analysis.

#### Histologic analysis of tumor tissue

Treated and untreated tumors were harvested from rats following euthanasia in accordance with institutionally approved protocols. Tumors were hemisected, fixed overnight in formalin, and dehydrated with 70% alcohol before paraffinization. 4-µm-thick sections were stained with hematoxylin and eosin (H&E) or Cibacron Brilliant Red-3BA (CBR-3BA) in accordance with previously described protocols.[35] Imaging was performed with Leica Aperio slide scanner microscope (Leica Microsystems, Wetzlar, Germany). Pieces of tumor tissue were prepared for electron microscopy via standard methods.[59] Sections were examined using a Talos L120C.

#### In vivo CT imaging

CT images were obtained in vivo using a MILabs μCT scanner (Utrecht, The Netherlands) with the following settings: peak voltage = 50 kV, current = 240 μA, step angle = 0.75°, and exposure time = 75 ms. Rats were scanned at 24-48 h post-embolization. Analysis of the images was performed using OsiriX.

#### Statistical analysis

All the experiments were performed independently in at least three replicates unless stated otherwise. In all figures, data points represent the mean, and error bars are the standard deviations or the standard error of mean, as specified. ANOVA with Bonferroni’s multiple comparison test was performed to establish statistical differences in anti-cancer effect of hydrogels drug loading *in vitro*. A Tukey’s multiple comparison test, with a single pooled variance was used to determine if there is a significant difference in attenuation between different AuNP loadings. P-values ≤ 0.05 were considered statistically significant. All statistical analyses were carried out using Graphpad Prism 8 software (San Diego, California).

## Acknowledgements

This work was supported by the NIH (R21-EB029556).

## Notes

### Competing Interest Statement

The authors have declared no competing interest.

## References

1. Bruix J, Gores GJ, Mazzaferro V. Hepatocellular carcinoma: clinical frontiers and perspectives. Gut. 2014; 63: 844–55.

2. Facciorusso A. Drug-eluting beads transarterial chemoembolization for hepatocellular carcinoma: Current state of the art. World J Gastroenterol. 2018; 24: 161–9.

3. Luz JHM, Luz PM, Martin HS, Gouveia HR, Levigard RB, Nogueira FD, et al. DEB TACE for Intermediate and advanced HCC - Initial Experience in a Brazilian Cancer Center. Cancer imaging. 2017; 17: 5-.

4. Marrero JA, Kulik LM, Sirlin CB, Zhu AX, Finn RS, Abecassis MM, et al. Diagnosis, Staging, and Management of Hepatocellular Carcinoma: 2018 Practice Guidance by the American Association for the Study of Liver Diseases. Hepatology (Baltimore, Md). 2018; 68: 723–50.

5. Balogh J, Victor D, 3rd, Asham EH, Burroughs SG, Boktour M, Saharia A, et al. Hepatocellular carcinoma: a review. J Hepatocell Carcinoma. 2016; 3: 41–53.

6. Jou JH, Muir AJ. Hepatocellular Carcinoma Surveillance. Clinical gastroenterology and hepatology : the official clinical practice journal of the American Gastroenterological Association. 2018; 16: 19–20.

7. Dimitroulis D, Damaskos C, Valsami S, Davakis S, Garmpis N, Spartalis E, et al. From diagnosis to treatment of hepatocellular carcinoma: An epidemic problem for both developed and developing world. World journal of gastroenterology. 2017; 23: 5282–94.

8. Raza A, Sood GK. Hepatocellular carcinoma review: current treatment, and evidence-based medicine. World journal of gastroenterology. 2014; 20: 4115–27.

9. West H, Jin JO. Transarterial Chemoembolization. JAMA Oncology. 2015; 1: 1178-.

10. White JA, Redden DT, Bryant MK, Dorn D, Saddekni S, Abdel Aal AK, et al. Predictors of repeat transarterial chemoembolization in the treatment of hepatocellular carcinoma. HPB : the official journal of the International Hepato Pancreato Biliary Association. 2014; 16: 1095–101.

11. Nguyen QV, Lym JS, Huynh CT, Kim BS, Jae HJ, Kim YI, et al. A novel sulfamethazine-based pH-sensitive copolymer for injectable radiopaque embolic hydrogels with potential application in hepatocellular carcinoma therapy. Polymer Chemistry. 2016; 7: 5805–18.

12. Kalva SP, Iqbal SI, Yeddula K, Blaszkowsky LS, Akbar A, Wicky S, et al. Transarterial chemoembolization with Doxorubicin-eluting microspheres for inoperable hepatocellular carcinoma. Gastrointestinal cancer research : GCR. 2011; 4: 2–8.

13. Ramsey DE, Kernagis LY, Soulen MC, Geschwind J-FH. Chemoembolization of hepatocellular carcinoma. J Vasc Interv Radiol. 2002; 13: S211–S21.

14. Kennedy AS, Nutting C, Coldwell D, Gaiser J, Drachenberg C. Pathologic response and microdosimetry of 90Y microspheres in man: Review of four explanted whole livers. Int J Radiat Oncol Biol Phys. 2004; 60: 1552–63.

15. Gaba RC, Yap FY, Martinez EM, Li Y, Guzman G, Parvinian A, et al. Transarterial sorafenib chemoembolization: Preliminary study of technical feasibility in a rabbit model. J Vasc Interv Radiol. 2013; 24: 744–50.

16. Parvinia A, Casadaban LC, Hauck ZZ, Van Breemen RB, Gaba RC. Pharmacokinetic study of conventional sorafenib chemoembolization in a rabbit VX2 liver tumor model. Diagnostic Interv Radiol. 2015; 21: 235–40.

17. Raoul JL, Heresbach D, Bretagne JF, Ferrer DB, Duvauferrier R, Bourguet P, et al. Chemoembolization of hepatocellular carcinomas a study of the biodistribution and pharmacokinetics of doxorubicin. Cancer. 1992; 70: 585–90.

18. Malagari K, Chatzimichael K, Alexopoulou E, Kelekis A, Hall B, Dourakis S, et al. Transarterial chemoembolization of unresectable hepatocellular carcinoma with drug eluting beads: results of an open-label study of 62 patients. Cardiovascular and interventional radiology. 2008; 31: 269–80.

19. Ashrafi K, Tang Y, Britton H, Domenge O, Blino D, Bushby AJ, et al. Characterization of a novel intrinsically radiopaque Drug-eluting Bead for image-guided therapy: DC Bead LUMI. Journal of controlled release : official journal of the Controlled Release Society. 2017; 250: 36–47.

20. Forster JC, Harriss-Phillips WM, Douglass MJ, Bezak E. A review of the development of tumor vasculature and its effects on the tumor microenvironment. Hypoxia (Auckland, NZ). 2017; 5: 21–32.

21. Vollherbst DF, Gockner T, Do T, Holzer K, Mogler C, Flechsig P, et al. Computed tomography and histopathological findings after embolization with inherently radiopaque 40μm-microspheres, standard 40μm-microspheres and iodized oil in a porcine liver model. PLoS ONE. 2018; 13: e0198911.

22. Gade TPF, Tucker E, Nakazawa MS, Hunt SJ, Wong W, Krock B, et al. Ischemia induces quiescence and autophagy dependence in hepatocellular carcinoma. Radiology. 2017; 283: 702–10.

23. Hatefi A, Amsden B. Biodegradable injectable in situ forming drug delivery systems. Journal of Controlled Release. 2002; 80: 9–28.

24. Gutowska A, Jeong B, Jasionowski M. Injectable gels for tissue engineering. The Anatomical record. 2001; 263: 342–9.

25. Ahmadi F, Oveisi Z, Samani SM, Amoozgar Z. Chitosan based hydrogels: characteristics and pharmaceutical applications. Research in pharmaceutical sciences. 2015; 10: 1–16.

26. McLaughlin SW, Cui Z, Starnes T, Laurencin CT, Kan HM, Wu Q, et al. Injectable thermogelling chitosan for the local delivery of bone morphogenetic protein. J Mater Sci Mater Med. 2012; 23: 2141–9.

27. Nair LS, Starnes T, Ko J-WK, Laurencin CT. Development of injectable thermogelling chitosan–inorganic phosphate solutions for biomedical applications. Biomacromolecules. 2007; 8: 3779–85.

28. Nair LS, Starnes T, Ko J-WK, Laurencin CT. Development of Injectable Thermogelling Chitosan–Inorganic Phosphate Solutions for Biomedical Applications. Biomacromolecules. 2007; 8: 3779–85.

29. Liu J, Yu M, Zhou C, Yang S, Ning X, Zheng J. Passive tumor targeting of renal-clearable luminescent gold nanoparticles: long tumor retention and fast normal tissue clearance. J Am Chem Soc. 2013; 135: 4978–81.

30. Soo Choi H, Liu W, Misra P, Tanaka E, Zimmer JP, Itty Ipe B, et al. Renal clearance of quantum dots. Nature Biotechnology. 2007; 25: 1165–70.

31. Gade TPF, Tucker E, Nakazawa MS, Hunt SJ, Wong W, Krock B, et al. Ischemia Induces Quiescence and Autophagy Dependence in Hepatocellular Carcinoma. Radiology. 2017; 283: 702–10.

32. Chandra VM, Wilkins LR, Brautigan DL. Animal Models of Hepatocellular Carcinoma for Local-Regional Intraarterial Therapies. Radiol Imaging Cancer. 2022; 4: e210098.

33. Kurma K, Manches O, Chuffart F, Sturm N, Gharzeddine K, Zhang J, et al. DEN-Induced Rat Model Reproduces Key Features of Human Hepatocellular Carcinoma. Cancers. 2021; 13: 4981.

34. Nishino M, Jackman DM, Hatabu H, Yeap BY, Cioffredi LA, Yap JT, et al. New Response Evaluation Criteria in Solid Tumors (RECIST) guidelines for advanced non-small cell lung cancer: comparison with original RECIST and impact on assessment of tumor response to targeted therapy. AJR Am J Roentgenol. 2010; 195: W221–8.

35. Rossomacha E, Hoemanni CD, Shive MS. Simple Methods for Staining Chitosan in Biotechnological Applications. J Histotechnol. 2004; 27: 31–6.

36. Chen Z, Zhao M, Liu K, Wan Y, Li X, Feng G. Novel chitosan hydrogel formed by ethylene glycol chitosan, 1,6-diisocyanatohexan and polyethylene glycol-400 for tissue engineering scaffold: in vitro and in vivo evaluation. J Mater Sci: Mater Med. 2014; 25: 1903–13.

37. Zhu J-L, Yu SW-K, Chow PK-H, Tong YW, Li J. Controlling injectability and in vivo stability of thermogelling copolymers for delivery of yttrium-90 through intra-tumoral injection for potential brachytherapy. Biomaterials. 2018; 180: 163–72.

38. Zhao X, Liu Y, Jia P, Cheng H, Wang C, Chen S, et al. Chitosan hydrogel-loaded MSC-derived extracellular vesicles promote skin rejuvenation by ameliorating the senescence of dermal fibroblasts. Stem Cell Research & Therapy. 2021; 12: 196.

39. Malik NS, Ahmad M, Minhas MU, Tulain R, Barkat K, Khalid I, et al. Chitosan/Xanthan Gum Based Hydrogels as Potential Carrier for an Antiviral Drug: Fabrication, Characterization, and Safety Evaluation. Front Chem. 2020; 8.

40. Hamedi H, Moradi S, Hudson SM, Tonelli AE. Chitosan based hydrogels and their applications for drug delivery in wound dressings: A review. Carbohydr Polym. 2018; 199: 445–60.

41. Pellá MCG, Lima-Tenório MK, Tenório-Neto ET, Guilherme MR, Muniz EC, Rubira AF. Chitosan-based hydrogels: From preparation to biomedical applications. Carbohydr Polym. 2018; 196: 233–45.

42. Ruel-Gariépy E, Shive M, Bichara A, Berrada M, Le Garrec D, Chenite A, et al. A thermosensitive chitosan-based hydrogel for the local delivery of paclitaxel. Eur J Pharm Biopharm. 2004; 57: 53–63.

43. Zhou HY, Zhang YP, Zhang WF, Chen XG. Biocompatibility and characteristics of injectable chitosan-based thermosensitive hydrogel for drug delivery. Carbohydr Polym. 2011; 83: 1643–51.

44. Wang QQ, Kong M, An Y, Liu Y, Li JJ, Zhou X, et al. Hydroxybutyl chitosan thermo-sensitive hydrogel: a potential drug delivery system. Journal of Materials Science. 2013; 48: 5614–23.

45. Zhang Z, Cen C, Qian K, Li H, Zhang X, Zhang H, et al. Assessment of the embolization effect of temperature-sensitive p(N-isopropylacrylamide-co-butyl methylacrylate) nanogels in the rabbit renal artery by CT perfusion and confirmed by macroscopic examination. Scientific Reports. 2021; 11: 4826.

46. Salis A, Rassu G, Budai-Szűcs M, Benzoni I, Csányi E, Berkó S, et al. Development of thermosensitive chitosan/glicerophospate injectable in situ gelling solutions for potential application in intraoperative fluorescence imaging and local therapy of hepatocellular carcinoma: a preliminary study. Expert Opin Drug Deliv. 2015; 12: 1583–96.

47. Wang Q, He Y, Shen M, Huang L, Ding L, Hu J, et al. Precision Embolism: Biocompatible Temperature-Sensitive Hydrogels as Novel Embolic Materials for Both Mainstream and Peripheral Vessels. Advanced Functional Materials. 2021; 31: 2011170.

48. Lym JS, Nguyen QV, Ahn DW, Huynh CT, Jae HJ, Kim YI, et al. Sulfamethazine-based pH-sensitive hydrogels with potential application for transcatheter arterial chemoembolization therapy. Acta Biomater. 2016; 41: 253–63.

49. Si-Mohamed S, Cormode DP, Bar-Ness D, Sigovan M, Naha PC, Langlois J-B, et al. Evaluation of spectral photon counting computed tomography K-edge imaging for determination of gold nanoparticle biodistribution in vivo. Nanoscale. 2017; 9: 18246–57.

50. Cormode DP, Si-Mohamed S, Bar-Ness D, Sigovan M, Naha PC, Balegamire J, et al. Multicolor spectral photon-counting computed tomography: in vivo dual contrast imaging with a high count rate scanner. Scientific Reports. 2017; 7: 4784.

51. GRAS Notice No. GRN 000997. 2022.

52. Yan D, Li Y, Liu Y, Li N, Zhang X, Yan C. Antimicrobial Properties of Chitosan and Chitosan Derivatives in the Treatment of Enteric Infections. Molecules. 2021; 26: 7136.

53. Bouche M, Dong YC, Sheikh S, Taing K, Saxena D, Hsu JC, et al. A novel treatment for glioblastoma delivered by a radiation responsive and radiopaque hydrogel. ACS Biomater Sci Eng. 2021; 7: 3209–320.

54. Dong YC, Nieves LM, Hsu JC, Kumar A, Bouche M, Krishnan U, et al. A Novel Combination Treatment for Melanoma: FLASH Radiotherapy and Immunotherapy Delivered by a Radiopaque and Radiation Responsive Hydrogel. Chem Mater. 2023; 35: 9542–51.

55. Cheheltani R, Ezzibdeh RM, Chhour P, Pulaparthi K, Kim J, Jurcova M, et al. Tunable, biodegradable gold nanoparticles as contrast agents for computed tomography and photoacoustic imaging. Biomaterials. 2016; 102: 87–97.

56. McLaughlin SW, Cui Z, Starnes T, Laurencin CT, Kan HM, Wu Q, et al. Injectable thermogelling chitosan for the local delivery of bone morphogenetic protein. J Mater Sci Mater Med. 2012; 23: 2141–9.

57. Chen MH, Wang LL, Chung JJ, Kim Y-H, Atluri P, Burdick JA. Methods To assess shear-thinning hydrogels for application as injectable biomaterials. ACS Biomater Sci Eng. 2017; 3: 3146–60.

58. Thankam FG, Muthu J. Biosynthetic hydrogels--studies on chemical and physical characteristics on long-term cellular response for tissue engineering. J Biomed Mater Res A. 2014; 102: 2238–47.

59. Chhour P, Naha PC, O’Neill SM, Litt HI, Reilly MP, Ferrari VA, et al. Labeling monocytes with gold nanoparticles to track their recruitment in atherosclerosis with computed tomography. Biomaterials. 2016; 87: 93–103.

