## Supplemental Information for "Drug-Eluting, Radiopaque, Tumor-Casting Hydrogels for Endovascular Locoregional Therapy of Hepatocellular Carcinoma"

* Corresponding author

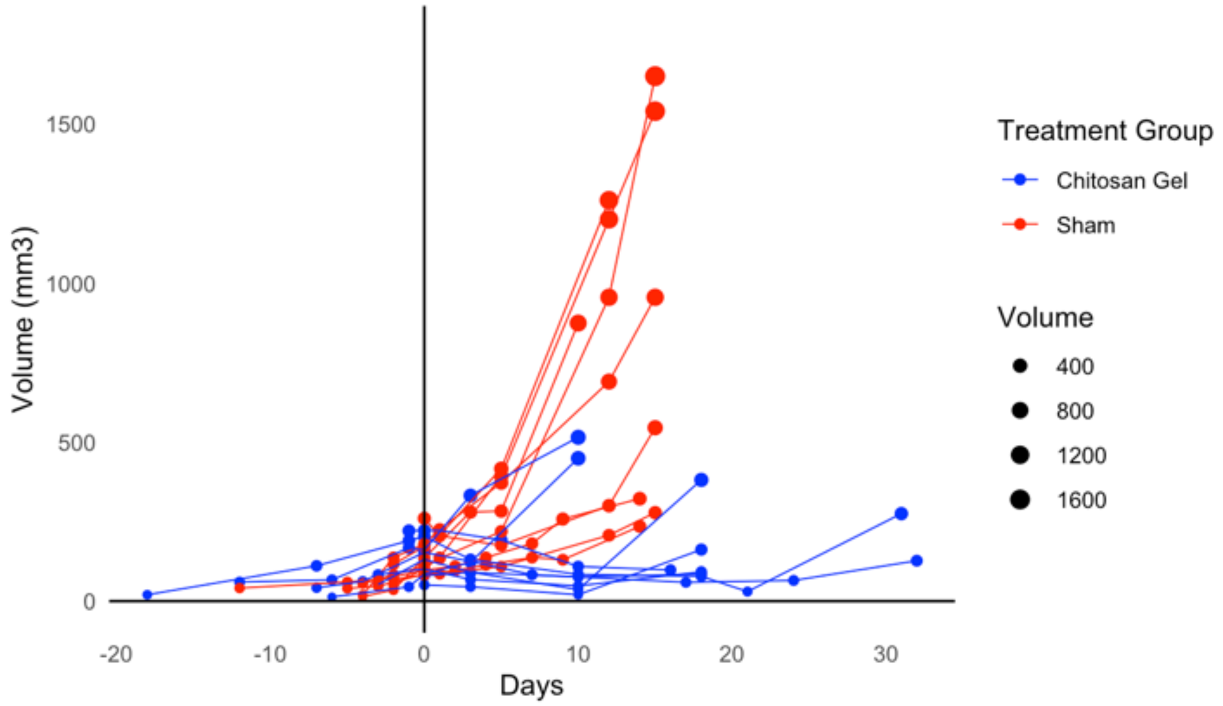


Figure S1. Comparison of the growth curves of tumors treated either with sham or chitosan gel.

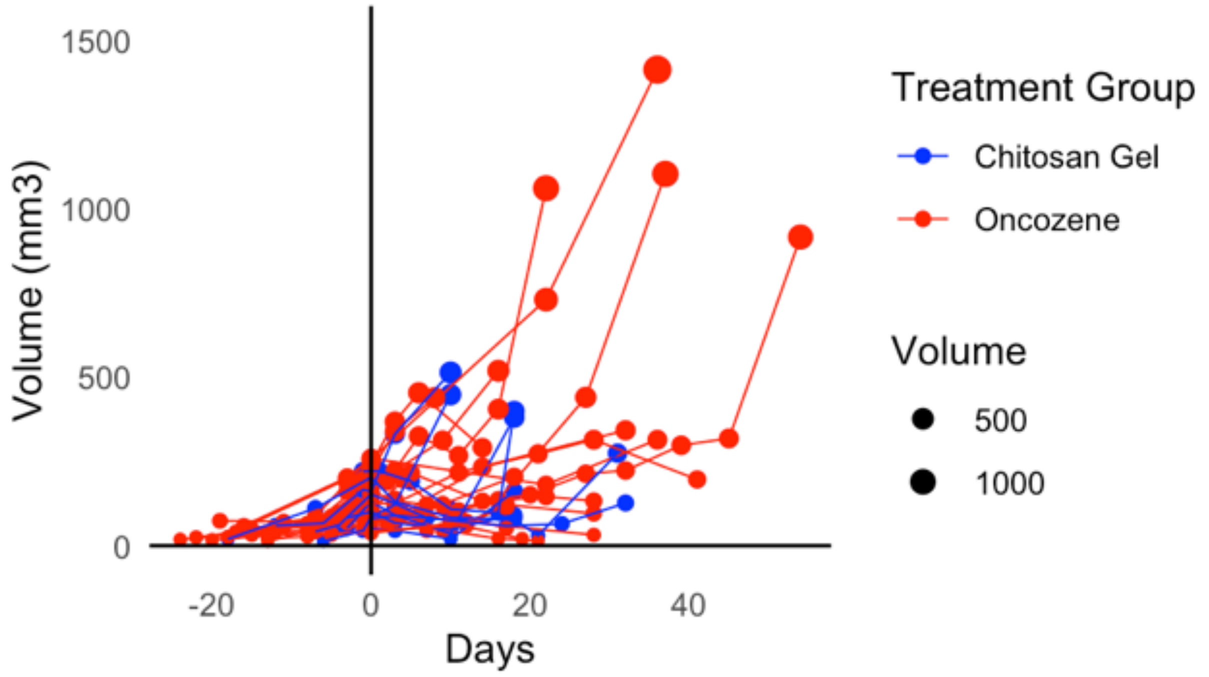


Figure S2. Comparison of the growth curves of tumors treated either with Oncozene beads or chitosan gel.
